# Tumor-associated macrophages confer resistance to chemotherapy (Trifluridine/Tipiracil) in digestive cancers by overexpressing Thymidine Phosphorylase

**DOI:** 10.1101/2024.08.19.608621

**Authors:** Marie Malier, Marie-Hélène Laverriere, Maxime Henry, Malika Yakoubi, Pascale Bellaud, Cécile Arellano, Anthony Sébillot, Fabienne Thomas, Véronique Josserand, Edouard Girard, Gael S Roth, Arnaud Millet

**Author notes:** **Corresponding author:** Arnaud Millet MD PhD, Team Mechanobiology, Immunity and Cancer, Institute for Advanced Biosciences, Allée des Alpes, 38700 La Tronche France.

## Abstract

Pyrimidine analogs are part of the first-line chemotherapy regimen for gastrointestinal cancers. Trifluridine combined with tipiracil, a specific thymidine phosphorylase inhibitor, in TAS-102 has recently emerged as a potential alternative in the face of primary or secondary chemoresistance to 5-fluorouracil. Despite its promise, in the current study, we report that macrophage-specific overexpression of thymidine phosphorylase results in macrophage-induced chemoresistance to TAS-102 that is insensitive to tipiracil inhibition. In addition, we demonstrate the human specificity of this mechanism, as mouse macrophages do not express significant levels of thymidine phosphorylase. To study the importance of macrophages in chemoresistance to trifluridine, we developed a humanized mouse model with tumor-implanted human macrophages and demonstrated their important role in treatment resistance to pyrimidine analogs. We also showed in human colorectal cancer that macrophages represent a major source of thymidine phosphorylase expression leading to chemoresistance.

**Significance:** Thymidine phosphorylase overexpression in TAMs confers chemoresistance to TAS-102 in digestive cancers.

## Introduction

Pyrimidine base analogs are an important component of chemotherapeutic regimens used in gastrointestinal cancers. 5-fluorouracil (5-FU), a tumor-inhibiting antimetabolite, is recognized as an effective molecule (Longley *et al*, 2003). However, many patients exhibit primary or secondary resistance to chemotherapeutic agents resulting in poor prognosis. Several mechanisms have been proposed to explain chemotherapeutic resistance (Tyner *et al*, 2022). Among these, the role of immune cells is of particular interest. We recently reported the importance of tumor-associated macrophages (TAMs) in resistance to 5-FU in colorectal cancers (Malier *et al*, 2021). We have shown that hypoxia-driven expression of dihydropyrimidine dehydrogenase (DPD) is a major source of resistance against 5-FU in a universal mechanism under the control of HIF-2α not limited to colorectal cancers. Facing resistance to a chemotherapeutic agent is a challenging clinical situation and can be circumvented by choosing another drug. Trifluridine (TFD) is another fluorinated pyrimidine analog that has shown strong potential against cancer cell growth (Fujiwara *et al*, 1970). This drug, combined with tipiracil, an inhibitor of the thymidine phosphorylase, under the name TAS-102, provided an increase in survival for patients with colorectal metastases in the RECOURSE trial (Mayer *et al*, 2015). This result justified the FDA and EMA approvals for its use in the treatment of patients with metastatic colorectal cancer who had previously been treated with 5-FU, oxaliplatin and irinotecan-based chemotherapy (Marcus *et al*, 2017). The interest of TAS-102 was then confirmed in association with bevacizumab, a vascular endothelial growth factor (VEGF)-inhibitor in the SUNLIGHT trial (Prager *et al*, 2023). The present study was designed to study the possibility to use TAS-102 as an alternative to 5-FU in macrophage-mediated chemo-resistance in digestive tumors.

## Results

### Macrophages confer chemoresistance to TAS-102 in a humanized tumor mice model

We recently showed that hypoxic macrophages confer chemoresistance to 5-FU in colorectal cancer (Malier *et al*, 2021). Indeed, 5-FU conditioning by hypoxic macrophages leads to cancer cells protection in a DPD-mediated mechanism as it is demonstrated by gimeracil blockade of DPD activity (Figure 1A). Surprisingly, macrophages confer also protection against TAS-102 in a DPD-independent mechanism as gimeracil is unable to revert this protection (Figure 1A). To further explore the role of macrophages in treatment resistance, we developed a humanized tumour model in NOG (NOD.Cg-*Prkdc^scid^ Il2rg^tm1Sug^*/JicTac) mice. The subcutaneous implantation of MIA-PaCa2, a human pancreatic cancer cell line, with human monocytes derived macrophages authorizes the follow-up of tumors with a human local myeloid compartment (Figure 1B). Macrophages do not interfere with tumor growth as it is illustrated by the follow-up of tumor volumes during the first 4 weeks after cancer cells implantation (Figure 1C). These macrophages display a phenotype similar to what is found in human tumors notably by expressing CD163 (Figure 1D), which is considered as a tumor-associated macrophages specific marker (Mantovani *et al*, 2021). Using this model, we sought to determine the relevance of macrophage-induced protection against TAS-102. Mice were treated with TAS-102 at 10 mg/kg twice a day (bid) orally during two cycles of 5 days starting 14 days after implantation of cells until day 28. Tumor size was measured until day 57 at which date mice were sacrificed (Figure 1E). We found that tumor associated macrophages confer resistance against TAS-102 as it is demonstrated by the tumor growth curves analysis after treatment (Figure 1F), tumor volume (Supplemental Figure 1) and tumor mass at end-point of the follow-up (Figure 1G).

**Figure 1.**
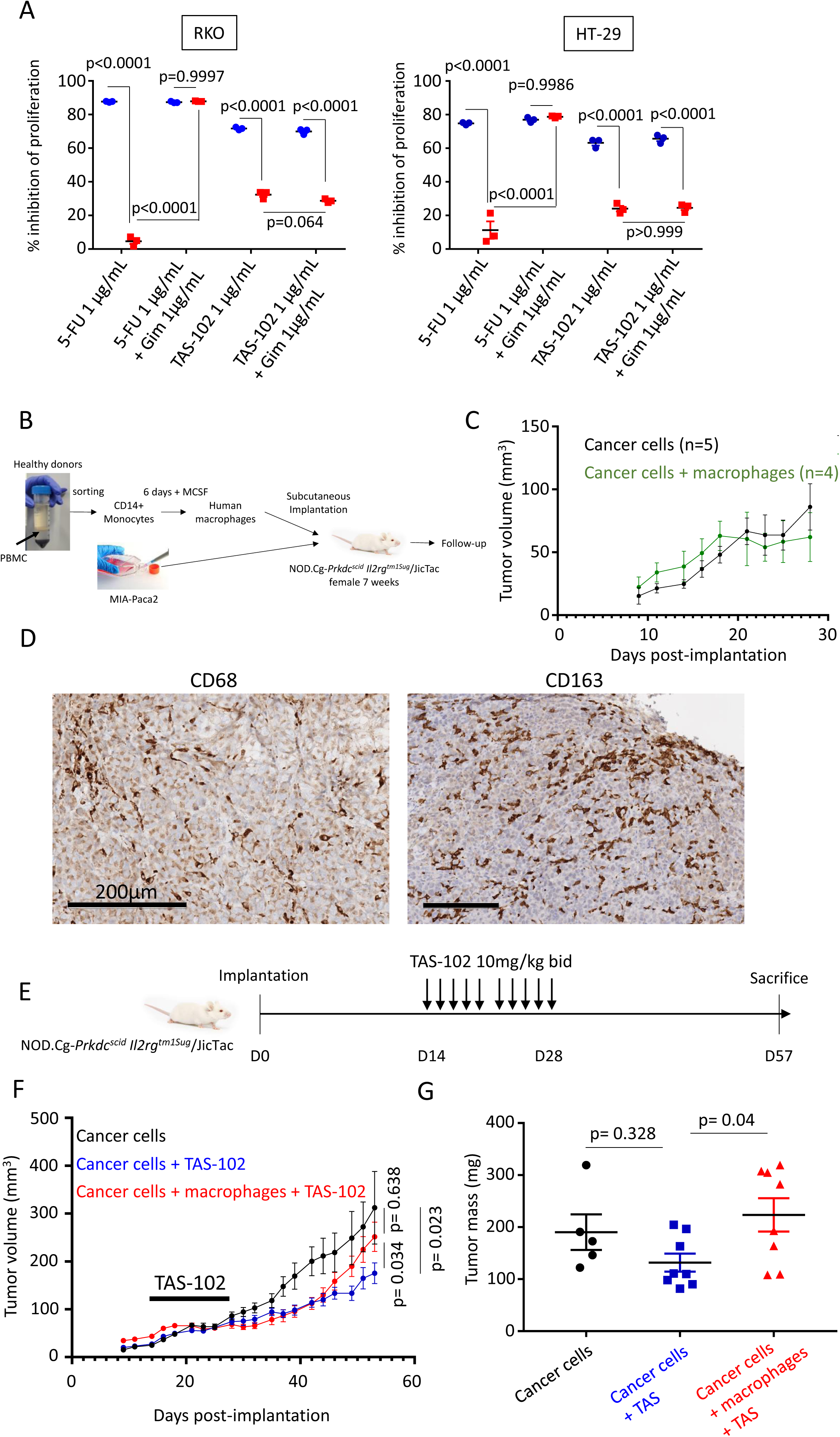
Macrophages confer chemoresistance to TAS-102 in a humanized tumor mice model. (A) RKO (Left panel) and HT-29 (Right panel) cultured with unconditioned medium (blue) or with macrophages conditioned medium (red) with 5-FU (1µg/mL), 5-FU (1µg/mL) + gimeracil (1µg/mL), TAS-102 (1µg/mL) and TAS-102 (1µg/mL) + gimeracil (1µg/mL) for 72h. Medium conditioning was performed during 24h. The mean +/- sem of proliferation inhibition (in percentage) are represented (n = 3). (B) Schematic of the *in vivo* model protocol. (C) Tumor growth analysis starting at day 9 after implantation until day 28 for cancer cells alone (black) and for cancer cells with macrophages (green). (D) Immunochemistry analysis of hCD68 and hCD163 expression in tumors from mice sacrificed at day 14 post-implantation. (E) Time-line of TAS-102 treatment for implanted NOG mice. (F) Tumor growth analysis for untreated tumors (black, n=5), tumors without human macrophages treated by TAS-102 (blue, n=8) and tumors with human macrophages treated by TAS-102 (red, n=8). (G) Tumor masses (mg) at day 57 for untreated tumors (n=5, black), tumors without human macrophages treated by TAS-102 (blue, n=8) and tumors with human macrophages treated by TAS-102 (red, n=8).

### Macrophages confer chemoresistance to trifluridine mediated by thymidine phosphorylase expression

TFD is the active pyrimidine analog composing TAS-102 (Figure 2A). To assess the mechanism involved in macrophage-driven protection against TAS-102 found in our in vivo model (Figure 1F), we cultured macrophages with cancer cells and exposed them to trifluridine. Macrophages were able to confer a strong resistance to the proliferation inhibition and apoptosis-induced by TFD (Figure 2B). The protection, provided by macrophages, is confirmed in a 3D culture of cancer cells (Figure 2C). We determined trifluridine IC50 on RKO or HT-29 cells and found the following values: 0.1 and 0.27 µg/mL respectively (Figure 2D). These IC50 increases above 2µg/mL when an indirect co-culture with macrophages is performed (Figure 2D). We further established the need for a direct contact between trifluridine and macrophages to obtain such protection. Indeed, the macrophage secretome alone is unable to provide such protection, demonstrating the prerequisite needed contact between trifluridine and macrophages to observe resistance to treatment (Figure 2E). Thymidine phosphorylase (TP) is an enzyme involved in pyrimidine base metabolism that catalyzes the conversion of thymidine to thymine. It has been shown that TP is able to facilitate the removal of the sugar moiety of trifluridine to yield a trifluoromethyluracil molecule (Figure 2F), which is inactive (Warfield & Reigan, 2022). HPLC quantification confirmed the complete disappearance of trifluridine (TFD) and the production of trifluromethyluracil (TFMU) when TFD is in direct contact with macrophages (Figure 1G). RNA interference resulting in TP decreased expression (Supplemental Figure 2) and activity (Figure 2H), confirmed that macrophage-driven chemoresistance against trifluridine is TP-dependent (Figure 2I).

**Figure 2.**
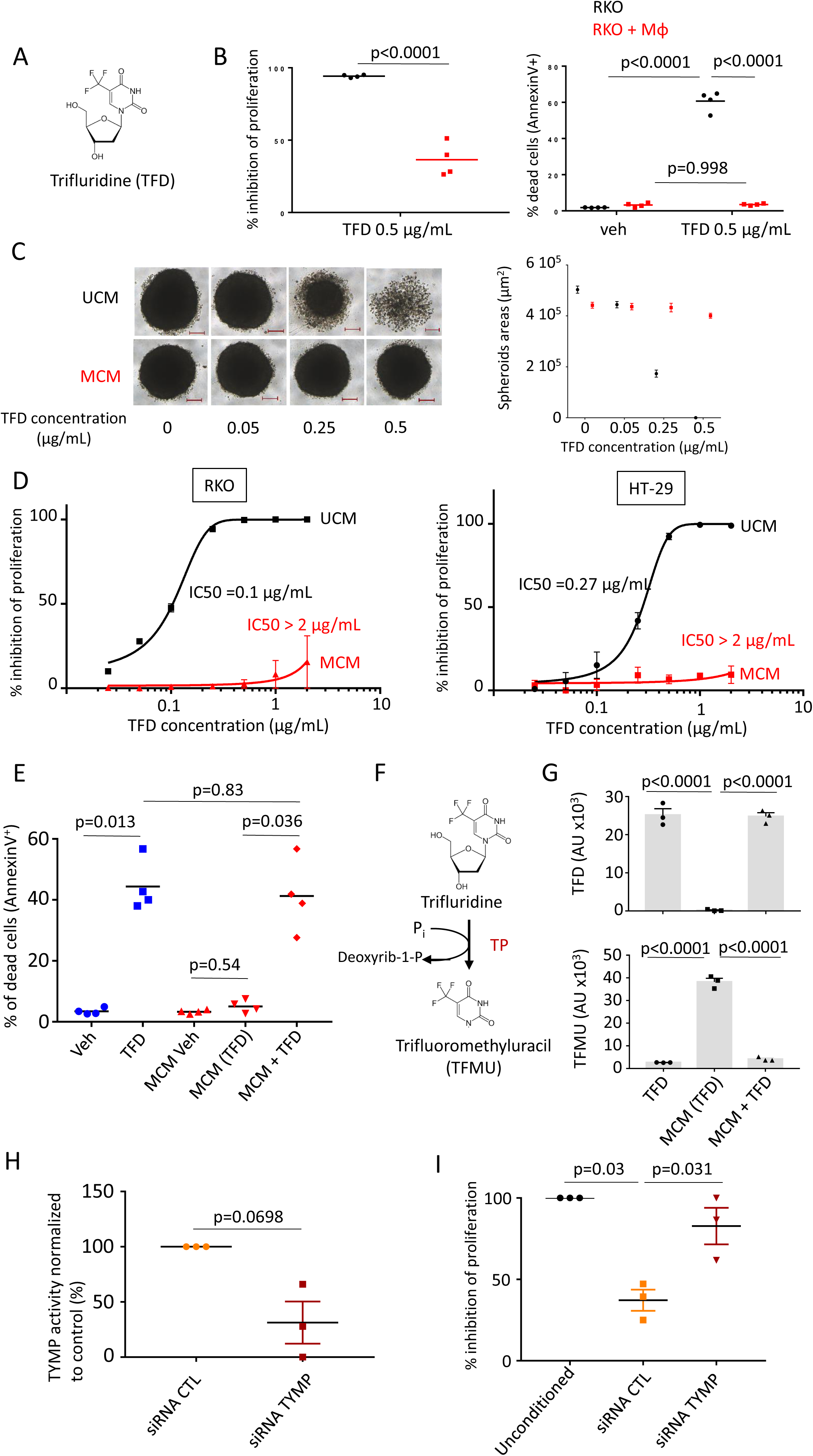
Macrophages confer chemoresistance to trifluridine mediated by thymidine phosphorylase expression. (A) Chemical structure of trifluridine (TFD). (B) Trifluridine activity at 0.5 µg/mL against RKO cells alone (black) or cocultured with macrophages (red) during an exposure time of 72h. Left panel represents the percentage of RKO inhibition of proliferation, right panel represents the percentage of RKO undergoing apoptosis evaluated by Annexin V staining (n=4). (C) HT-29 spheroids cultured with unconditioned medium (UCM, black, upper images) or with macrophages conditioned medium (MCM, red, lower images) at increasing TFD concentrations (scale bar = 200µm). The graph on the right represents the mean of spheroids areas in µm^2^ for each condition (n = 10 per condition), and the bar indicates the confidence interval at 95%. (D) RKO (Left panel) and HT-29 (Right panel) cultured with unconditioned medium (UCM, black) or with macrophages conditioned medium (MCM, red) containing an increasing dose of TFD from 0.025 to 2 µg/mL. The mean +/- sem values (%) of proliferation inhibition are represented (n = 4).(E) RKO cultured with unconditioned medium (UCM, blue), with macrophages conditioned medium (MCM (TFD), red), or with MCM in which TFD was added after the conditioning (MCM + TFD, red). The panel represents the percentage of RKO undergoing apoptosis (Annexin V positive staining) for 4 independent experiments. (F) TFD active conversion to trifluoromethyluracil (TFMU) and Deoxyrib-1-P under catalytic thymidine phosphorylase activity. (G) Dosage of trifluridine (TFD) and trifluoromethyluracil (TFMU) in medium with TFD for 24h, in macrophages conditioned medium with TFD for 24h noted as MCM(TFD) and in macrophages conditioned medium MCM with TFD for 24h added after 24h of conditioning noted as MCM + TFD (n=3). (H) TP activity in cytoplasmic extracts from macrophages treated by SiRNA scrambled (SiRNA CTL, orange) or siRNA directed against TYMP (SiRNA TYMP, dark red) (n=3). (I) RKO cultured for 48h with unconditioned medium (UCM, black) with TFD at 0.5 µg/mL, with conditioned medium containing TFD at 0.5 µg/mL during 6h by siRNA scrambled treated macrophages (siRNA CTL, orange) or by siRNA against TYMP treated macrophages (siRNA TYMP, dark red). The panel represents the percentage of RKO inhibition of proliferation (n=3).

### Tipiracil, a specific and potent TP inhibitor is unable to revert the macrophage-driven chemoresistance

Tipiracil (TPI), a specific inhibitor of TP, has been shown to increase trifluridine bioavailability in monkeys (Emura *et al*, 2005), leading to its use as a component of TAS-102, the trifluridine containing chemotherapy (Figure 3A). Surprisingly, macrophage-induced protection toward trifluridine is similar to TAS-102 (Figure 3B). Increasing dose of TPI slightly dampens that protection but is unable to restore trifluridine sensitivity even at 50µg/mL (Figure 3C & D). Of note the maximum plasmatic concentration of TPI in patients treated by TAS-102 is less than 100 ng/mL (Cleary *et al*, 2017). This absence of inhibition of macrophage’s TP activity contrasts with the potent inhibitory effect of TPI on TP activity in cytoplasmic extract from human macrophages for which the IC50 was measured at 8.1 nM (Figure 3E). We observed a similar activity of TPI against a human recombinant TP, showing a potent inhibitory power (Supplementary Figure 3). To appreciate the intracellular presence of TPI in macrophages, we exposed macrophage to various dose of TPI (0.5 and 50 µg/mL) during 24h and we analyzed TP activity, measured from cytoplasmic extracts. TPI is able to significantly inhibit TP activity only at the highest dose but not completely (Figure 3F), similarly to what we found previously (Figure 3C). Nevertheless, the TPI at 0.5 µg/mL (1.8 µM) which corresponds to the concentration used in TAS-102 (1.7 µM) when trifluridine is set at 1 µg/mL (3.4µM) is inefficient. This demonstrates that TPI is able to enter into the cell but not sufficiently to obtain a relevant inhibition. To assess if this finding is specific to macrophages, we performed the same experiment in BxPC3, a human pancreatic cancer cell line expressing TYMP. BxPC3 displays a similar pattern, advocating for a underappreciated inefficiency of TPI intracellular transport to block TP (Figure 3G). This result shows that TPI is able to penetrate intracellularly but the resulting intracellular TPI/TP molar ratio is insufficient to prevent TP-dependent degradation of trifluridine in human macrophages. To further rule out a potential degradation of TPI by macrophages, we analyzed the presence of 6-hydroxymethyluracile (6-HMU), a known degradation product of TPI (Peters, 2015), in macrophage supernatant. Indeed, 6-HMU represents less than 4% (in molar amount) of the total TPI excluding macrophage-induced degradation as a prominent mechanism explaining the inefficiency of TPI (Figure 3H).

**Figure 3.**
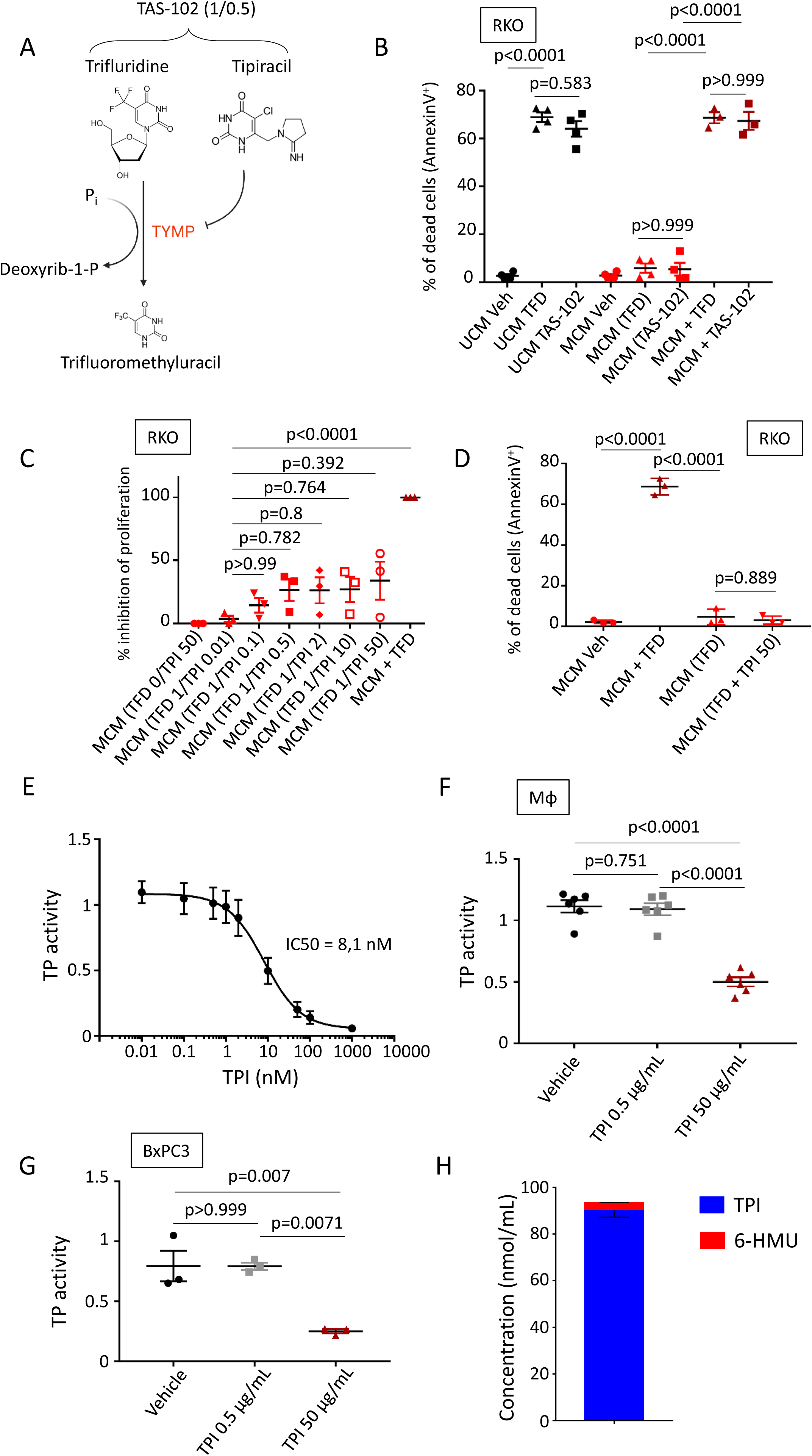
In physiological conditions, tipiracil is unable to restore chemosensitivity. (A) Chemical structure of TAS-102, composed of Trifluridine (TFD) and Tipiracil (TPI) in a 1/0.5 molar ratio. (B) Quantification of the apoptosis rate by flow cytometry (Annexin V positive cells) induced in RKO cells cultured with unconditioned medium (UCM, black) with vehicle (water), TFD or TAS-102, with macrophages conditioned medium with vehicle (MCM veh), TFD (MCM (TFD), red) or TAS-102 (MCM (TAS-102), red), or with MCM in which vehicle, TFD or TAS-102 were added after the conditioning (MCM + TFD, dark red). TFD was used at 1µg/mL and TAS-102 (TFD 1µg/mL and TPI 0.5µg/mL). (C) Proliferation rate evaluated for RKO cells exposed to macrophages conditioned medium containing TFD associated with increasing doses of TPI (µg/mL) (MCM (TFD/TPI), red) and with TFD (1µg/mL) added after the conditioning (MCM + TFD, dark red) (n=3). (D) Quantification of the apoptosis rate by flow cytometry (Annexin V positive cells) induced in RKO cells exposed to macrophages conditioned medium containing trifluridine at 1µg/mL (MCM (TFD) with or without TPI (50 µg/mL) in red and with TFD (1µg/mL) added after the conditioning MCM + TFD in dark red (n=3). (E) Response curve of TP enzyme activity from human macrophage cytoplasmic extracts (corresponding to 170 000 cells) with increasing dose of TPI. TYMP activity is measured by the conversion of trifluridine (TFD) into trifluoromethyluracil (TFMU) using absorbance quantification at 296 nm (n=4). (F) TP enzymatic activity from human macrophage cytoplasmic extracts of cells with prior exposuse to vehicle, TPI at 0.5µg/mL or 50 µg/mL for 24h. TP activity was measured by the conversion of trifluridine (TFD) into trifluoromethyluracil (TFMU) using absorbance quantification at 296 nm (n=6). (G) TP enzymatic activity from human BxPC3 cytoplasmic extracts of cells with prior exposuse to vehicle, TPI at 0.5µg/mL or 50 µg/mL for 24h. TYMP activity is measured by the conversion of trifluridine (TFD) into trifluoromethyluracil (TFMU) using absorbance quantification at 296 nm (n=3). (H) Quantification of 6-hydroxymethyluridine (6-HMU), the main degradation product of TPI, in the supernatant of macrophages exposed to TPI (50 µg/mL) during 24h (n=3).

### Macrophages represent the main source of TYMP expression in normal tissues in humans

Using single cell RNAseq data from normal donors (from the Human Protein Atlas data), we can assess the level of expression of various cell types in normal digestive tissues. The pattern of expression of *TYMP* for cells present in digestive tissues reveals the predominance of expression in the monocytic cell lineage (Figure 4A). Indeed, the hierarchical clustering of genes and cell types shows that *TYMP* is one of the specific gene to identify monocytic derived cells in normal tissues (Figure 4B & 4C). This pattern is confirmed in another dataset (Human Cell Landscape) showing that monocyte/macrophage lineage is the main source of tissue *TYMP* (Figure 4D). The predominance of TP monocytic expression compared to other immune cells (T and B lymphocytes) is confirmed by immunoblot (Supplementary Figure 4A). Interestingly, the protein level expression of TP is higher in monocyte derived macrophages than monocytes themselves (Supplementary Figure 4B). Interestingly, *TYMP* pattern expression in mice is not similar to humans. Indeed, mice monocytes express less *tymp* mRNA than hepatocytes or lymphocytes contrary to what is observed in humans (Figure 4E). The comparison between the enzymatic activity of TP from human monocyte derived macrophages and mice bone marrow derived macrophages (C57BL/6 and BALB/c) from cytoplasmic extracts confirmed the discrepancy between mice and humans showing no significant activity in mice macrophages (Figure 4F). These results indicate that mice macrophages should not be used as a model to assess tumor associated macrophages involvement in trifluridine treatment resistance.

**Figure 4.**
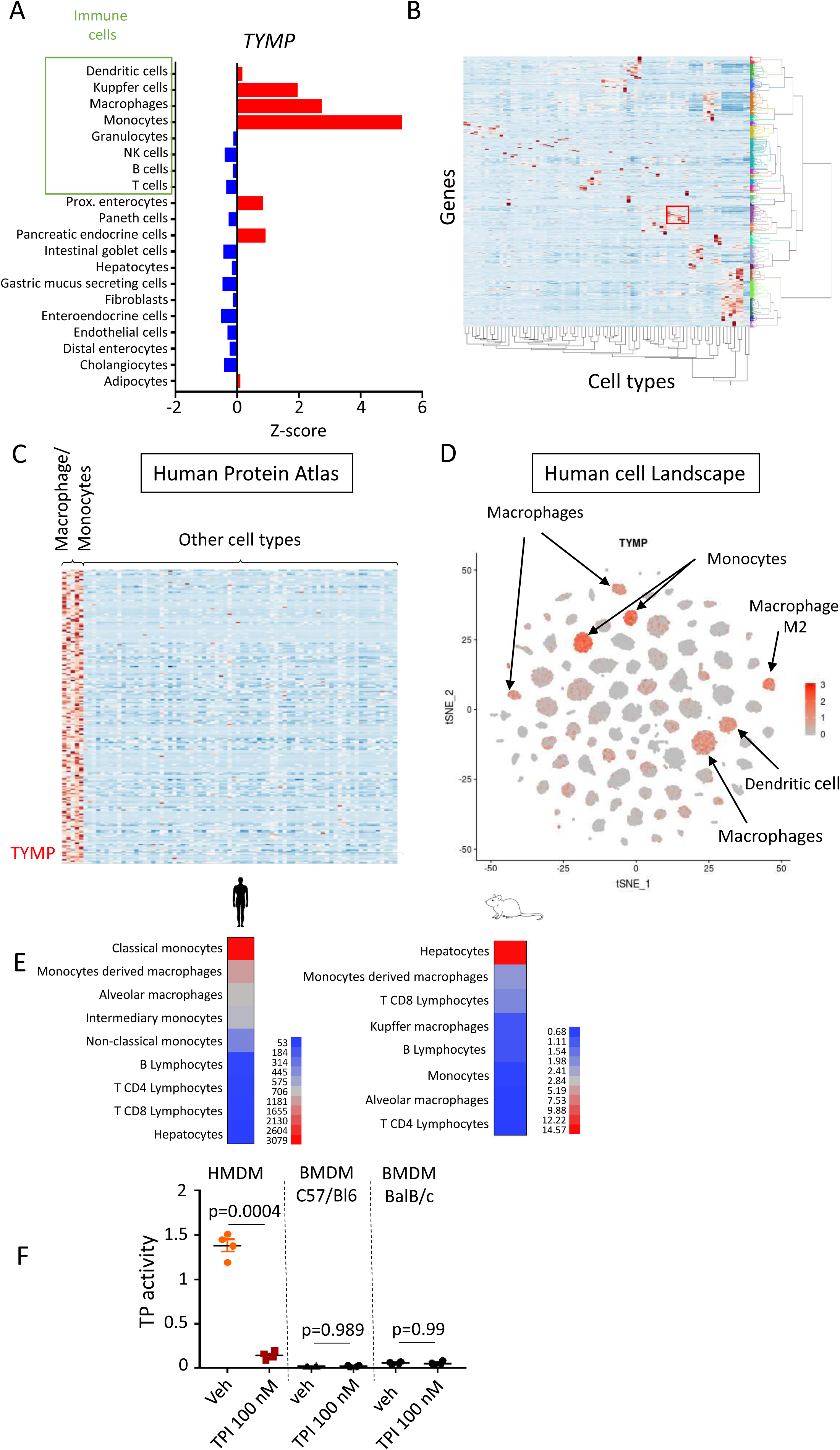
Macrophages are a major source of TP expression in normal tissue. (A) Single cell RNAseq expression of *TYMP* in cell types encountered in human digestive tissues from the Human Protein Atlas. (B) Heat-map of genes clustered according to their expression in the 79 cell types identified. The red square identifies the group of myeloid cells comprising macrophages, monocytes, dendritic cells and kuppfer cells. (C) Heat map of the various genes of the myeloid cluster defined in Figure 4B from the data of the Human Protein Atlas showing *TYMP* as one of the gene defining this cluster. (D) t-SNE heat map of *TYMP* expression from single cell RNAseq analysis of the Human Cell Landscape. (E) mRNA levels of expression of *TYMP* determined by microarray in human and mice in various cell types. (F) TP enzyme activity from cytoplasmic extracts (corresponding to 200000 cells) of human monocyte derived macrophages and mice bone marrow derived macrophages (C57Bl/6 and BalB/C). TP activity was measured by the conversion of trifluridine (TFD) into trifluoromethyluracil (TFMU) using absorbance quantification at 296 nm (n=4 in each group). TPI at 100 nM was used to inhibit the enzymatic activity due to TP.

### Macrophages represent the main source of TYMP expression in digestive tumor tissues in humans

In the context of colorectal cancer, single cell RNAseq analysis (GSE178341) shows a similar pattern of expression for TYMP, which is mainly expressed in the monocytic cell lineage (Figure 5A & 5B). This pattern could be extended to other digestive cancers. Indeed, THP1 monocytic cells express high TYMP levels compared to digestive cancer cell lines (Figure 5C). Moreover, THP1 differentiated macrophages display a higher TP protein expression than un-differentiated THP1-monocytes (supplementary Figure 5A). As human monocyte derived macrophages present a TP protein level expression superior to THP1 macrophages, the resulting pattern confirmed the predominance of macrophages as a source of TP and trifluridine resistance in various types of solid digestive cancers (pancreatic and colorectal) (Figure 5D). We have shown that macrophages confer resistance to TAS-102 in a humanized mouse model (Figure 1F). In this model, macrophages are the only source of TP expression in tumors as it was demonstrated by the absence of TP or CD163 expression in tumors lacking human macrophages (Supplementary Figure 5B). Mouse macrophages were detected at the tumor periphery according to their CD68 expression profile (CD68^+^CD163^-^TP^-^) (Supplementary Figure 5B). Quantification of human macrophage populations (TP^+^ (CD163^+^ or CD68^+^)), in this model, shows that they represent in average 16.6% of the cells in tumors (Figure 5E). To further explore the role of TP expression in tumors, we quantified its expression in non-tumor and tumor areas of patients with colic adenocarcinoma. TP expression was assessed by immunofluorescence and macrophages were identified by their expression of CD68 and/or CD163. In non-tumor tissues, macrophages were mainly found in the *lamina propria* (Figure 5F). In tumor tissues, macrophages are mainly found in infiltrative areas (figure 5G). Quantification of the cell populations shows that tumors contain a high proportion of myeloid cells, which can enter the monocyte/macrophage lineage thanks to their expression of CD68 or CD163. Among these cells we have identified macrophages expressing TP (TP+) and monocyte-like cells that do not express significant levels of TP (but are CD68^+^ or CD163^+^). These monocyte-like cells seems to be more abundant in non-tumor tissues than in tumor tissues contrary to macrophages which are more prominent in tumor tissues (Figure 5H). We also found that macrophages represent around 40% of the TP^+^ cell compartment in colon tissues (both non-tumor and tumor), demonstrating their key role in TP expression in tumor tissues (Figure 5I).

**Figure 5.**
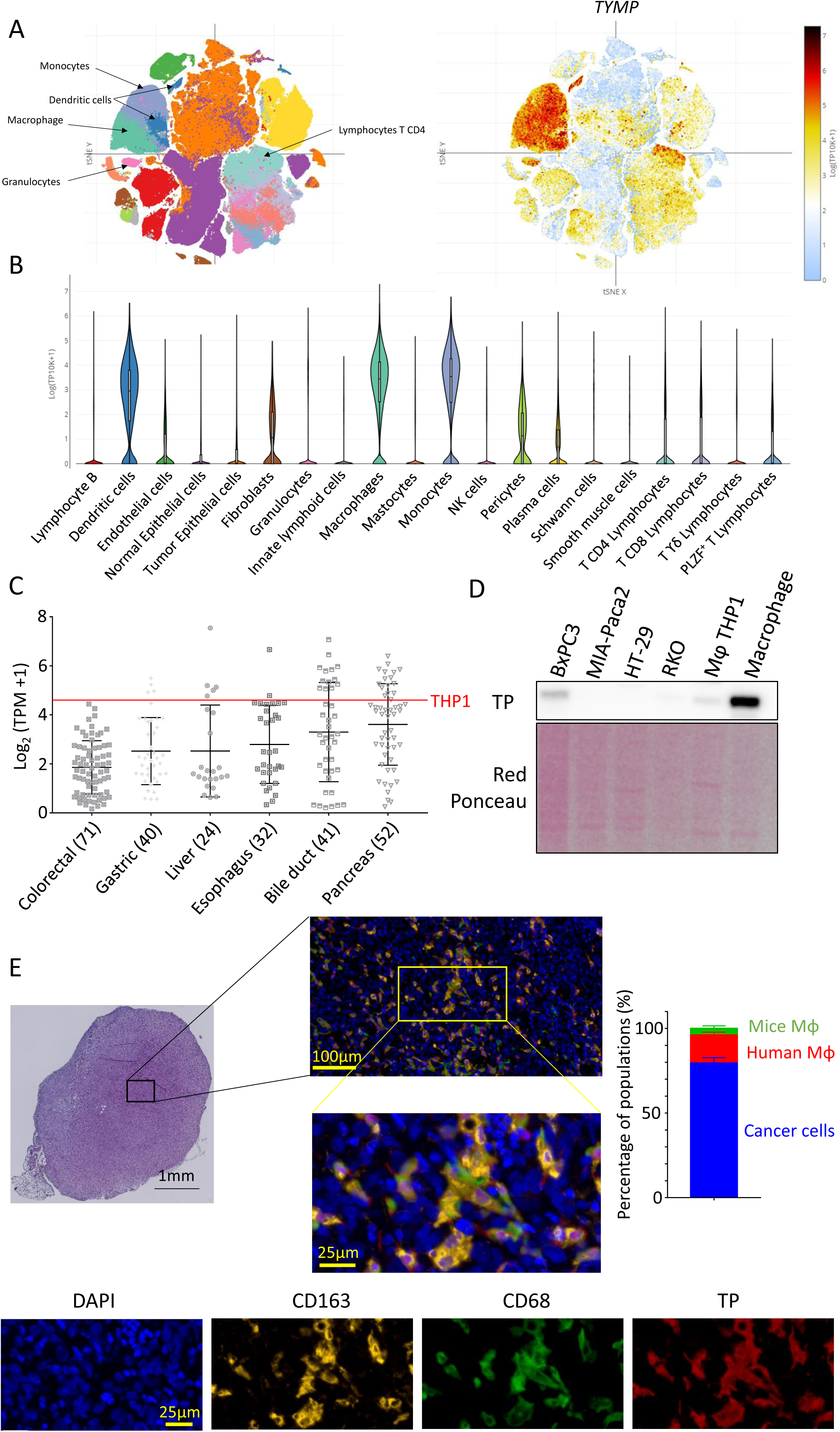
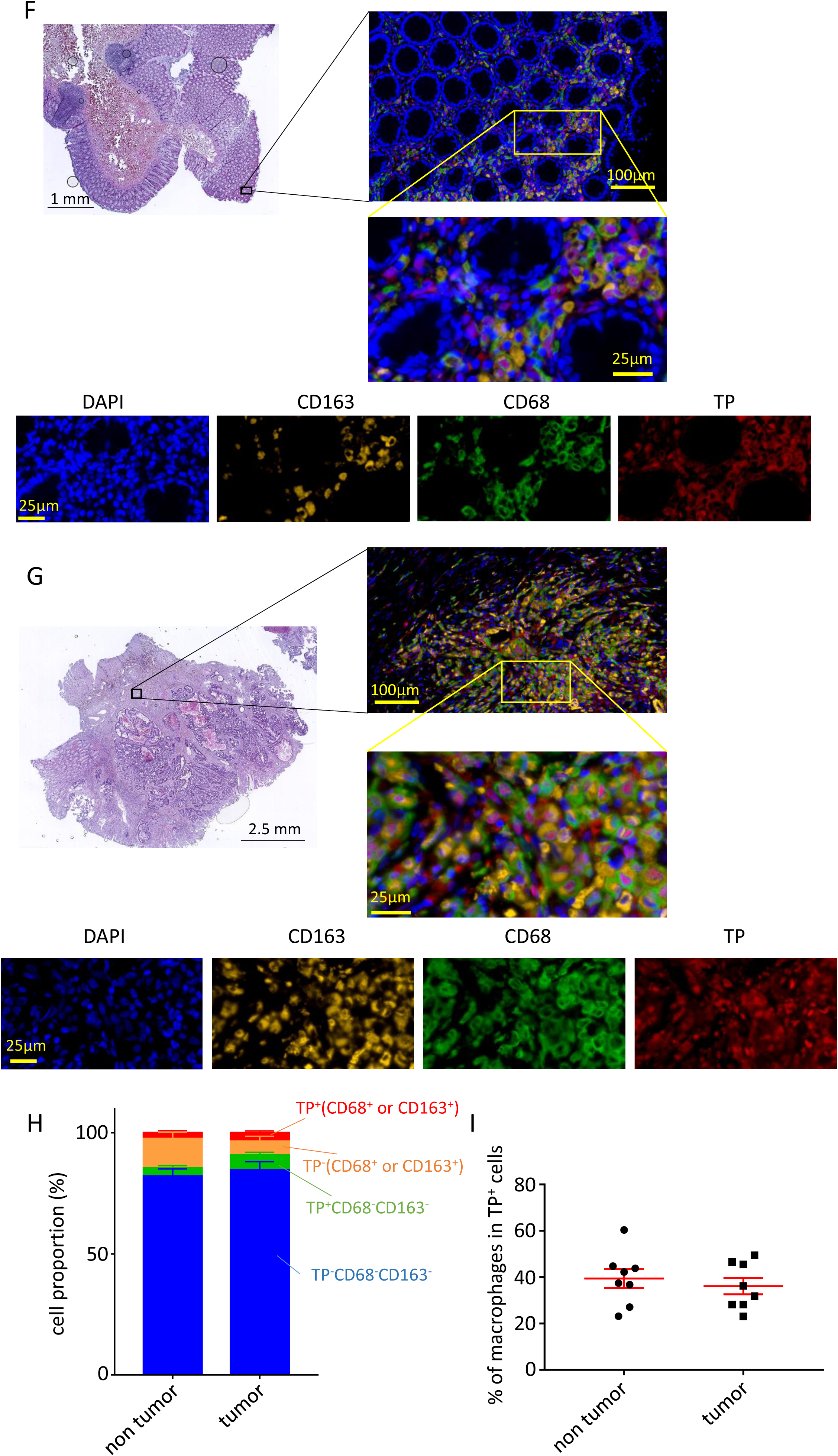
Macrophages are a major source of TP expression in tumor tissue. (A) tSNE heat map of single cell RNAseq with cell identification (left panel) and *TYMP* expression in the various cell clusters (right panel) in human colorectal cancers. (B) Violin plot of *TYMP* expression in various cell types from colorectal tumors. (C) RNAseq analysis of *TYMP* expression in various cancer cell lines from the CCLE (Cancer Cell Line Encyclopedia). THP1 mRNA level for *TYMP* is represented by the red line. (D) Immunoblot analysis of TP expression in cancer cell lines, THP1 macrophages and Human monocyte derived macrophages. Protein extract were deposited corresponding to 15,000 cells. (E) Immunofluorescence analysis of CD68, CD163, TP expression, in our mice model, in tumors 4 weeks after implantation corresponding to the end of the chemotherapy treatment. Nuclei were stained with DAPI. Quantification of cells populations were performed using the following classification: cancer cells (CD68^-^CD163^-^TP^-^), mice macrophages (CD68^+^CD163^-^TP^-^) and human macrophages (TP^+^). (F) Immunofluorescence analysis of TP, CD68, CD163 expression in human colon non-tumor tissues. Nuclei were stained with DAPI. (G) Immunofluorescence analysis of TP, CD68, CD163 expression in human colon tumor tissues. Nuclei were stained with DAPI. (H) Quantification of various cells proportions from human non-tumor and tumor tissues according to their TP, CD68 and CD163 expression pattern. (I) Quantification of macrophages in the TP+ cell compartment in human non-tumor and tumor tissues (n=8 in each group).

## Discussion

The immune system is increasingly recognized as a key element in understanding the mechanisms of tumor interactions with its surrounding healthy tissue and as a provider of new therapeutic targets. The tumor immune microenvironment (TIME) is composed of different types of immune cells. However, TAMs typically represent the quantitatively largest population of immune cells found in solid tumors. TAMs are involved in tumor growth, immune evasion, neoangiogenesis and treatment resistance. A large number of studies have reported an increased chemo-sensitivity to paclitaxel, doxorubicin, etoposide or gemcitabine when macrophages are depleted from the tumor in mice models (Ruffell & Coussens, 2015; Mitchem *et al*, 2013; Shree *et al*, 2011). Mechanisms involving macrophage-induced chemoresistance usually rely on the secretion of soluble factors by macrophages, such as pyrimidine nucleosides (deoxycytidine), which inhibit gemcitabine induction of apoptosis in pancreatic ductal adenocarcinoma (Halbrook *et al*, 2019). In colorectal cancer, the involvement of macrophages in chemoresistance has also been suggested based on *in vitro* and *in vivo* studies. The proposed mechanisms are diverse but also involve secreted factors influencing cancer cells response to the chemotherapeutic agent such as interleukin-6 (Yin *et al*, 2017).

5-FU is a widely used chemotherapeutic agent, particularly in gastrointestinal cancers. Its mechanism of action is complex, resulting in thymidine synthase dysfunction along with misincorporation of fluorodeoxyuridine (fdUTP) into RNA and DNA leading to cell proliferation blockade and death. Despite its potential efficacy, resistance to 5-FU is a frequent difficult clinical challenge. Many mechanisms have been proposed to explain the emergence of 5-FU resistance. One of the main explanations is based on the inactivation of the drug by DPD. This enzyme catalyzes the first step in the pyrimidine catabolic pathway. Many cancer cell types exhibit high levels of DPD expression mediated by miRNAs and kinases that promote DPD overexpression in cancer cells (Verma *et al*, 2022), except in colorectal cancers where DPD is downregulated mainly by negative epigenetic control (Malier *et al*, 2021; Wu *et al*, 2016). Over the past ten years, many strategies have been developed to address this problem using DPD inhibitors or DPD downregulators without success (Verma *et al*, 2022). Recently, we have shown that macrophages are a major source of 5-FU resistance in colorectal cancer due to the oxygen-mediated control of DPD expression in these cells (Malier *et al*, 2021). Because macrophages are one of the most abundant non-cancerous cell population in colorectal cancer, many patients are at risk for resistance. One way to circumvent 5-FU resistance is to use another drug such as TFD, a pyrimidine analog (Mayer *et al*, 2015; Prager *et al*, 2023). The triphosphate form of TFD is incorporated into DNA resulting in its cytotoxic effects (Tanaka *et al*, 2014). It appears that TFD is rapidly degraded by TP, justifying the association of TPI, a specific TP inhibitor. This compound, TPI, has been shown to increase the bioavailability of trifluridine in monkeys (Emura *et al*, 2005). The presence of TPI, in TAS-102, is expected to prevent TP-mediated TFD degradation in the tumor microenvironment. However, in the present study, we show that macrophages confer a strong resistance to TFD (Figures 2B-2E) in a TP-dependent mechanism (Figures 2H & 2I). The protection conferred by macrophages appears to be effective, as we report a complete protection against a challenge with 2µg/mL of TFD. Of note the maximum plasmatic concentration of TFD in humans suffering from solid cancers, was reported to be 2.155 µg/mL after a single dose of TAS-102 (Cleary *et al*, 2017). Surprisingly, this macrophage-mediated resistance to TFD is extended to TAS-102 *in vivo* (Figure 1F & 1G) and *in vitro* (Figure 3C & 3D) where TPI is inefficient in blocking TP activity at physiological concentrations, of note the maximum plasmatic concentration of TPI measured is less than 70 ng/mL after a single dose of TAS-102 (Cleary *et al*, 2017). We also show that TPI is inefficient to inhibit TP activity in BxPC3, a pancreatic cancer cell line that expresses high levels of TP (Figure 3G), supporting a poor intracellular TPI/TP ratio regardless of cell type. This is in accordance with a previous study that found TPI unable to increase TFD sensitivity in cancer cell lines expressing TP (Temmink *et al*, 2005). To further explain the inefficiency of TPI, we examined the expression level of TP in human macrophages. We found that macrophages are one of the major cell sources of TP in normal and cancer tissues (Figures 4 & 5). Interestingly, we found that this finding appears to be specific to humans, as mouse macrophages do not exhibit a similar TP activity or expression (Figures 4 E & F). Another surprising result is the relative discrepancy between mRNA level of *TYMP* in monocytes and macrophages resulting in an increased protein expression in macrophages (Supplemental Figure 4B). A potential mechanism is due to an intron retention in *TYMP* mRNA in monocytes released during differentiation toward macrophages (Green *et al*, 2020). These results reinforce our previous observation that the pyrimidine metabolism pathway is particularly high in human macrophages resulting in their key role in pyrimidine analogs resistance in digestive cancers. However, the expression pattern of human macrophages with respect to these two key enzymes, DPD and TP, has not been clearly elucidated. Accordingly, macrophages appear to be a relevant biomarker to predict treatment efficacy and represent a valuable therapeutic target to reverse chemoresistance, as we recently reported using a tyrosine kinase-specific inhibitor against CSF1R which is able to downregulate DPD expression in TAMs (Gharzeddine *et al*, 2024).

## Material and methods

### Resources

All references of products used in this study are available in the supplemental material and method file.

### Human Tissue Samples

Approval for the use of clinical samples and information was obtained from each patient and the Research Ethics Committee Ouest IV (CPP 55/21_3). Written informed consent was obtained prior to patient participation. This study is part of a registered clinical trial: NCT05038358 (clinicaltrial.gov). Clinical information are reported in the supplemental table S1.

### Cell Culture

RKO, HT-29, MIAPacCa2, BxPC3 and THP-1 were purchased from ATCC or ECACC and maintained in RPMI (Gibco) supplemented with 10% FBS (Gibco) at 37°C. Spheroids were produced using 500 HT-29 cells in 96-wells plate Nunclon Sphera (Life Technologies). All cells were routinely tested for mycoplasma contamination using MycoAlert detection kit (Lonza). All cells were used within 5 years of receipt.

### Human Macrophages Differentiation from Monocytes

Human blood samples from healthy de-identified donors were obtained from EFS (French National Blood Service) under an approved protocol (CODECOH DC-2022–5248). The donors gave signed consent for their blood to be used in this study. Monocytes were isolated from the leukoreduction system chambers of healthy EFS donors by differential centrifugation (Histopaque 1077, Sigma) to obtain PBMCs. CD14^+^ microbeads (Miltenyi Biotec) were used to select monocytes according to the manufacturer’s instructions. Monocytes were plated at 750 000 cells / well in 12-well plates in RPMI (Life Technologies) supplemented with 10% SAB (Sigma), 10 mM HEPES (Life Technologies), MEM non-essential amino acids (Life Technologies) (hereafter called macrophage medium) and 25 ng/mL M-CSF (Miltenyi Biotec). Differentiation was achieved after 6 days of culture. CD3^+^ T lymphocytes and CD20^+^ B lymphocytes were sorted from PBMCs using CD3^+^ and CD20^+^ microbeads (Miltenyi Biotec) according to the manufacturer’s instructions.

### Animals

7 weeks-old NOD.Cg-*Prkdc^scid^ Il2rg^tm1Sug^*/JicTac (NOG) female mice were purchased from Taconic. Animals were housed at the Institute for Advanced Biosciences Animal Core Facility (Grenoble, France), Agreement number (D 38 516 10001), under specific pathogen–free conditions, in a temperature-controlled environment with a 12-hour light/dark cycle and ad libitum access to water and food. Animal husbandry and procedures were performed according to the recommendations of the Direction des Services Vétérinaires, Ministry of Agriculture of France, according to the Council Directive 2010/63/EU of the European Communities, and according to recommendations for the health monitoring from the Federation of European Laboratory Animal Science Associations. Protocols involving animals were reviewed by the local ethics committee “Comité d’Ethique pour l’Expérimentation Animale n°12, Cometh-Grenoble” and approved by the Ministry of Research under the approval number APAFIS# 33137-2021110411585349 v2.

### Macrophage Conditioned Medium

Monocytes were plated at 750 000 cells / well in 12-well plates. Fully differentiated human macrophages were used to condition medium with vehicle (water) or trifluridine (TFD) at 1 µg/mL for 24 hours, unless otherwise stated. Target cells (HT-29 or RKO) were plated at 25 000 cells / well in 24-well plates 24 hours prior to exposure to conditioned medium. Target cells were incubated with unconditioned medium (UCM) or macrophage conditioned medium (MCM) for 48 hours. They were then counted using an Attune flow cytometer (Life Technologies), and the inhibition of proliferation was determined. Apoptosis was assessed using Annexin V – FITC staining and analyzed by flow cytometry.

### Co-cultures experiments

Monocytes were plated at 500 000 cells / well in 12-well plates. On day 6 and 9, the medium was changed. On day 12, RKO were plated with the macrophages at 50 000 cells / well. After 24 hours, the medium was changed and replaced by control macrophage medium (vehicle) or trifluridine (TFD) at 0.5 µg/mL. After 72 hours, RKOs were harvested, counted and stained with Annexin V for apoptosis. Macrophages that could have been retrieved with RKOs were identified and excluded according to their SIRPα expression (RKO are SIRPα^-^ and macrophages are SIRPα^+^).

### RNAi

Fully differentiated macrophages were transfected with siTYMP (L-009281-00-0005, Dharmacon) at a final concentration of 50 nM using Lipofectamine (RNAiMAx, Life Technologies) on day 6 and 12, with medium changes on day 9 and 16. They were used on day 19.

### Measurement of TP activity

For cytoplasmic extracts, cells were counted, then lyzed in 20 mM Tris-Hcl, 150 mM NaCl, 0.1% Triton, 2 mM EDTA, 1 mM AEBSF, 400 µM leupeptin, 1 µM pepstatin A, 10% glycerol and 1 mM DTT lysis buffer. TP activity assessment was based on trifluridine (TFD) conversion to trifluoromethyluracil (TFMU). Briefly, TFD was put at 333 µM in 96-wells UVstar plates (Greiner), human recombinant TP or cellular extracts from BxPC-3, human macrophages or mouse BMDM were added to TFD. After 6 hours at 37°C, the reactions were stopped by adding 3 µL of NaOH 10 M, and the absorbance at 296 nm was read on a CLARIOstar (BMG Labtech)(Kaspar *et al*, 2019). The quantification was made by subtracting the 296 nm absorbance at t=0 to the 296 nm absorbance after 6 hours.

### *In vivo* experiments

5x10^6^ human macrophages and 10^6^ Mia-PaCa2 human cancer cells were implanted subcutaneously into NOG mice. Mice were distributed into 4 groups: MIA-Paca2 treated with vehicle (n=5), MIA-PaCa2 treated with TAS-102 (n=8), Macrophages + MIA-Paca2 treated by vehicle (n=4) and Macrophages + MIA-PaCa2 treated by TAS102 (n=8). Two weeks after implantation, mice were treated with TAS-102 at 10mg/kg orally bid for two consecutive weeks (5 days). Tumor size was measured every two working days using a digital caliper along two orthogonal axes (∥ was the largest one and ⊥ was the smallest one) and tumor volume was determined using the following formula 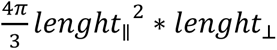. Mice were sacrificed 8 weeks after tumor implantation and tumors were harvested, weighed, then fixed in paraformaldehyde and embedded in paraffin for histological analysis.

### Mouse and human mRNA *TYMP* expression analysis

Genevestigator 7.5.0 (https://genevestigator.com/gv/) is a search engine that summarizes data sets into metaprofiles. GENEVESTIGATOR® integrates manually curated and quality controlled gene expression data from public repositories (Hruz *et al*, 2008). In this study, the Condition Tools Search was used to obtain *TYMP* mRNA levels obtained from human (*Homo sapiens*) and mouse (*Mus musculus*) in different anatomical parts. The mean of the logarithmic expression level obtained from AFFIMETRIX expression microarrays was used to generate a cell plot from selected results related to macrophages and monocytes. The lowest and the highest expression results in both series (human and mouse) were used to evaluate the expression level in the data set. The cell plot was generated using the JMP software (SAS).

### Single Cell RNAseq analysis

Single cell RNAseq analysis for healthy tissues was performed from a collection of scRNAseq datasets available from the Human Protein Atlas (www.humancellatlas.org). The single cell RNA sequencing datasets were based on a meta-analysis of the single cell RNA sequencing literature and single cell databases including healthy human tissues (Karlsson *et al*, 2021). To avoid technical bias and to ensure that the single cell dataset can best represents the corresponding tissue, the following data selection criteria were applied: (1) single cell transcriptomic datasets were limited to those based on the Chromium single cell gene expression platform from 10X Genomics (version 2 or 3); (2) single cell RNA sequencing was performed on single cell suspensions from tissues without pre-enrichment of cell types; (3) only studies with >4,000 cells and 20 million reads were included, (4) only datasets whose pseudo-bulk transcriptomic expression profile is highly correlated with the transcriptomic expression profile of the corresponding HPA tissue bulk sample were included. Exceptions were made for eye (∼12.6 million reads), rectum (2,638 cells) and heart muscle (plate-based scRNA-seq) to include different cell types in the analysis. Data extracted from this database were processed using JMP SAS software to generate heat maps and histograms. Single cell RNA-seq analysis for tumors was performed on open data from the Broad Institute under the GEO reference GSE178341 using the Global tSNE visualization tool (https://singlecell.broadinstitute.org/single_cell). Briefly, 371 223 cells from 28 mismatch repair-proficient (MMRp) and 34 mismatch repair-deficient (MMRd) colorectal tumors were transcriptionally profiled. Analysis of 88 cell subsets and their 204 associated gene programs was performed (Pelka *et al*, 2021).

### Immunofluorescence analysis

Paraffin-embedded tissue was sectioned at 4 µm, mounted on Adhesion slides (TOMO (ROCHE) and dried at 58°C for 12 hours. Immunofluorescence staining was performed on the Discovery ultra-automated IHC stainer, using the Discovery Rhodamine kit, the Discovery CY5 kit, the Discovery FAM kit (VENTANA). After deparaffination with VENTANA Discovery Wash Solution at 75 °C for 8 minutes, antigen retrieval was performed by using Tris-based buffer solution CC1 at 95°C to 100°C for 40 minutes. Endogenous peroxidase was blocked with Disc Inhibitor for 12 minutes at 37°C. After rinsing with reaction buffer, the slides were incubated with an 1/500e dilution of the first primary antibody (TP) for 60 minutes at 37°C. After rinsing, a secondary antibody, a goat anti-Rabbit HRP, was used to amplify the signal and incubated for 16 min and the discovery rhodamine kit incubated for 8 minutes. After denaturation with VENTANA solution CC2 at 100°C for 8 minutes, and Disc Inhibitor at 37°C for 8 minutes. Slides were incubated with a 1/100e dilution of second primary antibody (anti-CD68) at 37°C for 60 minutes. After rinsing, signal enhancement was performed using a secondary antibody, goat anti-Mouse HRP, and incubated for 16 min and the discovery CY5 kit incubated for 8 minutes. This was followed by denaturation with VENTANA Solution CC2 at 100°C for 8 minutes and Disc Inhibitor at 37°C for 8 minutes. Slides were incubated with a 1/200e dilution of the third primary antibody (anti-CD163) at 37°C for 60 minutes. After rinsing, signal enhancement was performed with a secondary antibody, goat anti-Rabbit HRP, incubated for 16 min and the discovery kit FAM incubated for 8 minutes. Slides were then counterstained with DAPI for 4 minutes and rinsed. After removal from the instrument, the slides were manually rinsed and coverslipped. Immunofluorescence staining was observed using a slide scanner (Hamamatsu NanoZoomer S60). HES staining was performed on an automated multistainer (LEICA ST 5020). Image analysis was performed using Qupath (v0.5.1)(Bankhead *et al*, 2017). Cell detection of objects was first performed using DAPI staining to identify nuclei then a classifier for CD68, CD163 and TP were trained and quantification performed using a composite classifier applied to annotated regions of interest. Measurements were then used to quantify proportions of various cell types in non-tumor and tumor tissues.

### TFMU, TFD, TPI and 6-HMU measurement

Assays were performed with a UPLC-MSMS, TQD (triple quadrupole, Waters) equipped with MassLynx software and using the following ionization settings: capillary voltage: 3Kv, cone voltage: 45 V, source temperature: 150°C, desolvation temperature: 500°C, gas flow rate: 50 L/H (cone) and 1000 L/H (desolvation), collision gas flow rate: 0.15 ml/min. Quantification of TFD; TPI and 6-HMU were performed in ESI positive mode at the following specific transitions (MRM mode, quantification transitions are indicated in bold): TFD : 297.0939>117.0397 ; 297.0939>98.949 and 297.0939>181.0152, TPI : 243.0251>183.0228, 243.0251>119.042 ; 243.0251>155.0006; 6-HMU : 142,9420>98.0104, 142,9420>81.9796 and 142,9420>97.9667. TFMU quantification was performed in ESI negative mode: 178.8549>42.082. Chromatographic separation was performed with an Acquity PREMIER UPLC HSS 1.8 m 2.1x150mm column (Waters), maintained at 40°C, at a flow rate of 0.5 ml/min, with a gradient between an aqueous phase A (0.1% formic acid in water) and an organic phase B (0.1% formic acid in acetonitrile) evolving as follow : T0-1min: 100%A, 2.5-4min: 5%A/95%B, 4.2min-7.5 min: 100%A. Calibration solutions were prepared by spiking RPMI media with TFD and TFMU, TPI and 6-HMU at concentrations ranging from 1 to 10000 ng/ml for TFD, tipiracil and 6-HMU and 0. 5-625 ng/mL for TFMU. [2-13C, 15N2] dihydrouracil (90 ng/mL) was used as internal standard (IS). Extraction procedure is performed as follows: 500 µl of cell supernatant or calibration points is spiked with 50µl of IS solution, then 600mg of ammonium sulfate is added to each solution and vortexed for 1 minute. Four mL of an Isopropanol/Ethyl acetate mixture (15:85,v/v) is dispensed into each tube, shaken with a rotary shaker for 15 minutes and then centrifuged at 3500g at 4°C for 15 minutes. The organic phase is then evaporated to dryness in a dry bath at 37°C under a stream of air. The residue is taken up with 120 µL of water containing 0.1% formic acid and vortexed for 5 minutes. Solutions are filtered and transferred to vials for analysis.

### Statistical analysis

Statistics were performed using Graph Pad Prism 7 (Graph Pad Software Inc). Data are presented as the mean ± sem. When two groups were compared, a paired or unpaired two-tailed student’s t-test was used when appropriate. When more than two groups were compared, a one-way ANOVA analysis with Sidak’s multiple comparison test on paired data was used when appropriate. Tumor growth curves were analyzed by two-ways ANOVA using the open-access TumorGrowth software (Enot *et al*, 2018). The likelihood of the data according to a null hypothesis (p-value) is presented in the figures. The number of independent experiments used to generate the data is shown in each legend’s figure.

## Authors’ contributions

A. **M. Malier**: Investigation, writing and editing. **MH. Laverierre**: Investigation (histology analysis). **M Henry**: Investigation (animal handling). **M. Yakoubi**: Investigation. **P Bellaud**: Investigation. **C. Arellano**: Investigation. **A Sébillot**: Investigation. **F Thomas**: Investigation. **V Josserand**: Investigation (animal handling and ethical issues). **E. Girard**: Investigation (patient recruitment). **G. Roth**: Investigation (patient recruitment). **A. Millet**: Conceptualization, resources, data curation, formal analysis, supervision, funding acquisition, validation, investigation, visualization, methodology, writing–original draft, project administration, writing–review and editing.

## Acknowledgments

A. Millet is supported by the ATIP/Avenir program (Inserm and La ligue nationale contre le cancer). This work is supported by the ERiCAN program of Fondation MSD-Avenir (Reference DS-2018-0015). We thank Bénédicte Demoustier for her technical help at the beginning of this project.

## Supplemental Figures

**Supplemental Figure 1.**
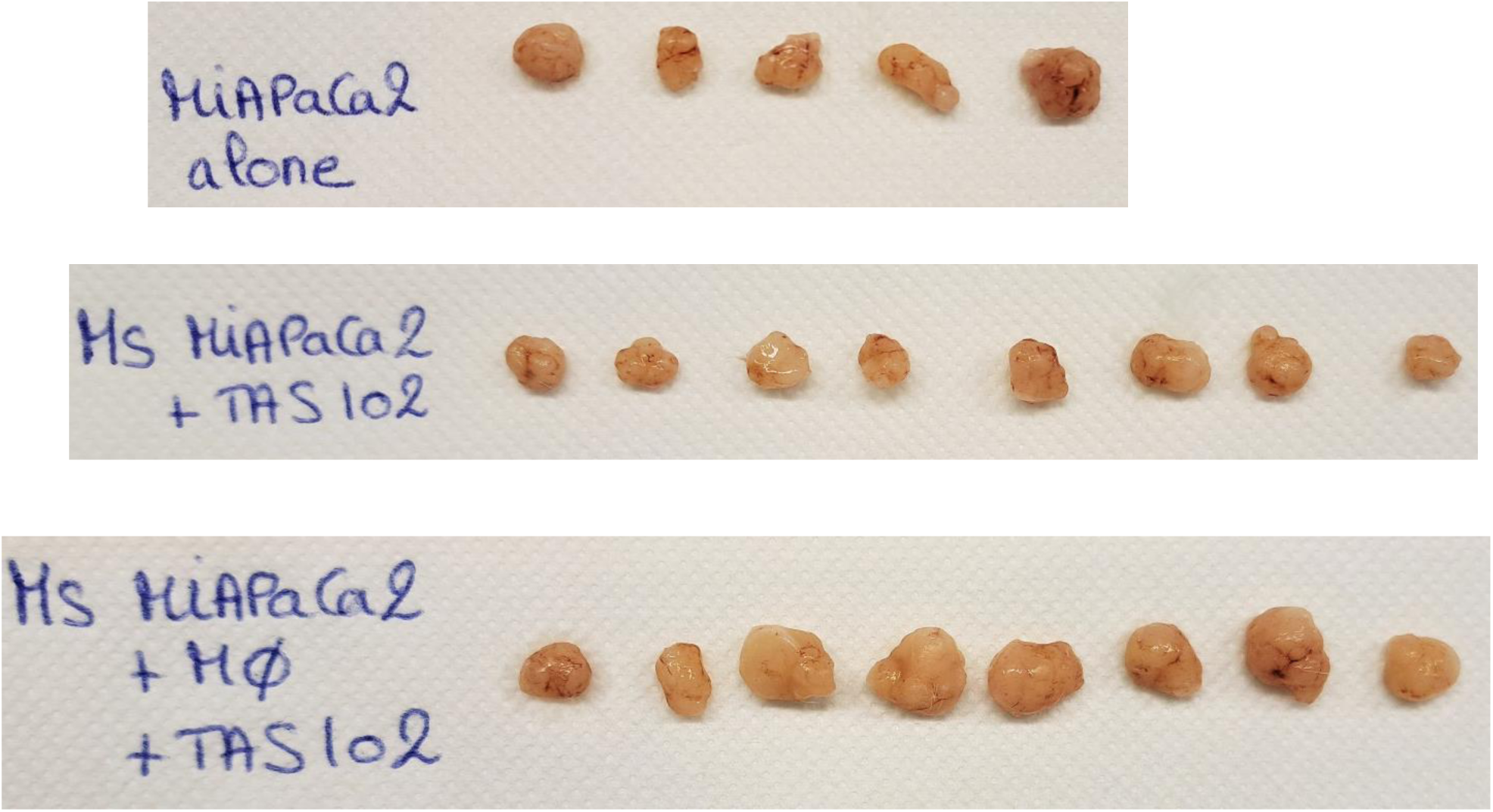
Pictures of tumors extracted from mice at the end of the follow-up at day 57 post-implantation.

**Supplemental Figure 2.**
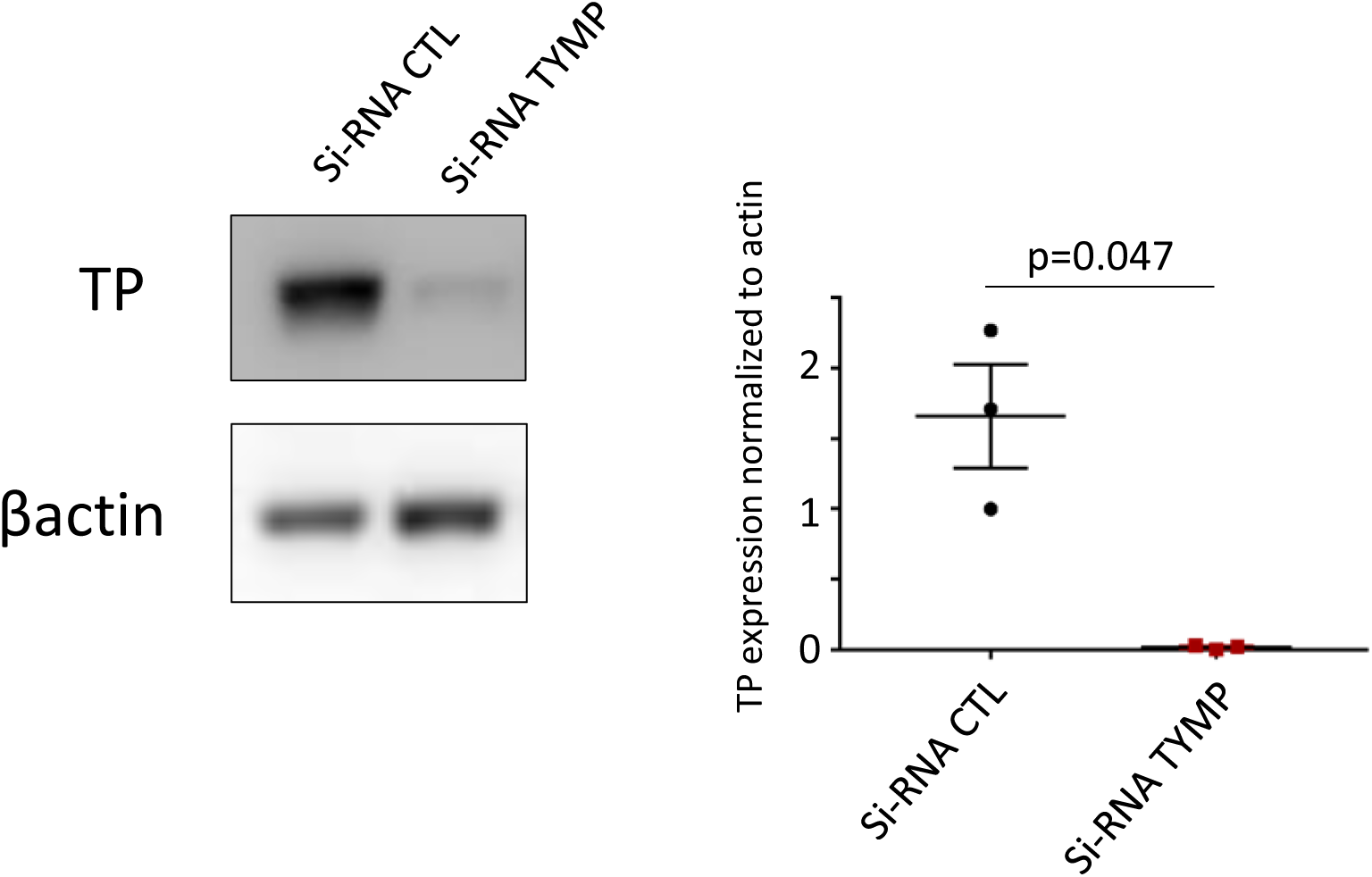
Immunoblot analysis of TP expression in macrophages treated by siRNA against TYMP. Representative of 3 independent human donors.

**Supplemental Figure 3.**
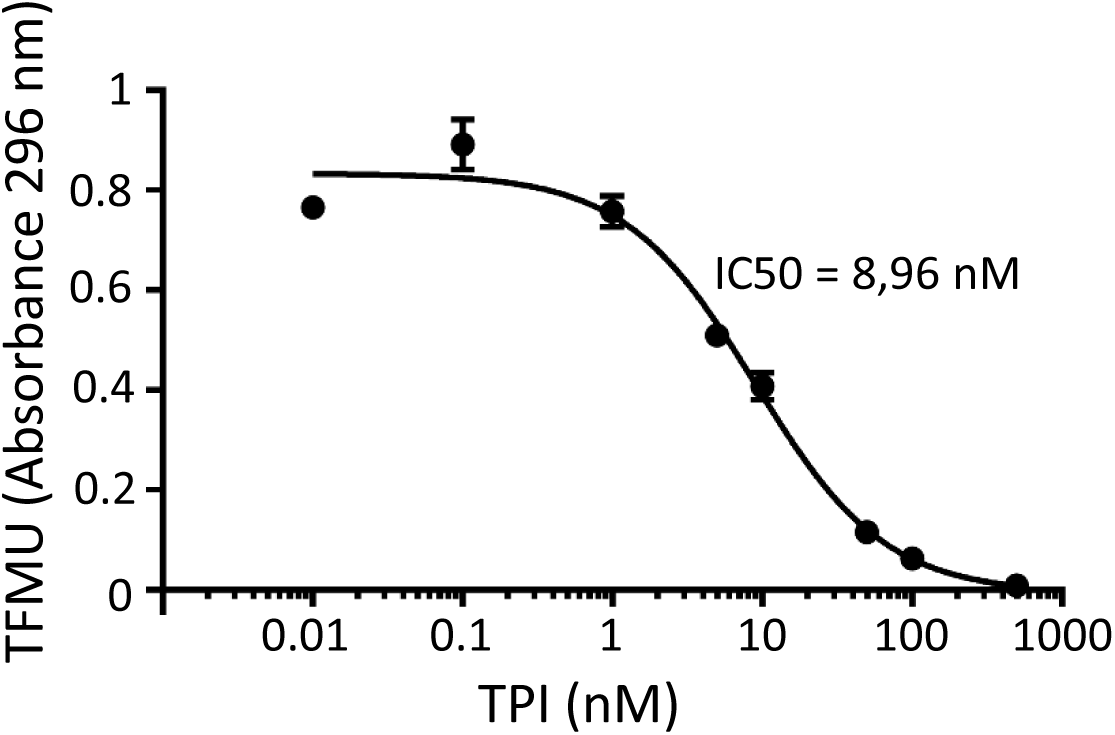
Response curve of human recombinant TP (100 ng) enzyme activity to increasing dose of TPI. TP activity was measured by the conversion of trifluridine (TFD) into trifluoromethyluracil (TFMU) using absorbance quantification at 296 nm (n=4).

**Supplemental Figure 4.**
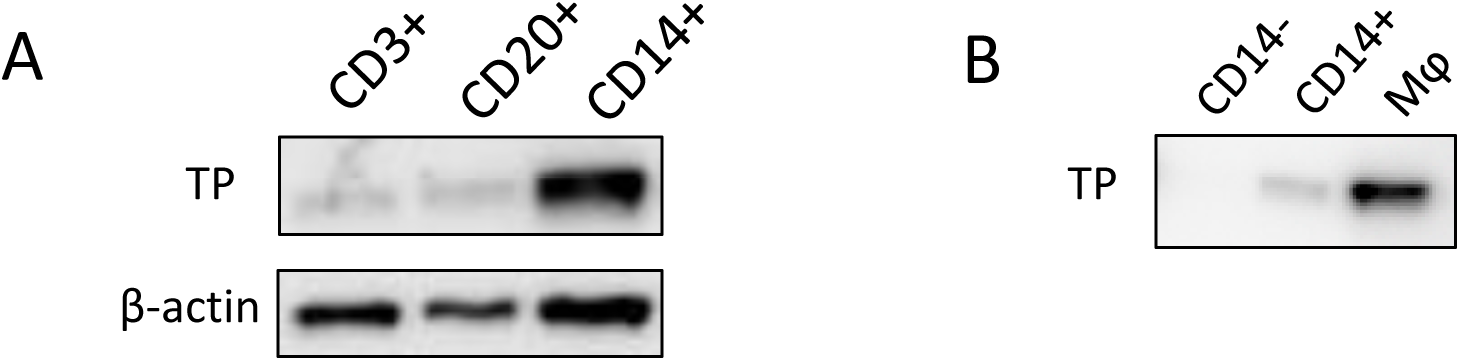
(A) Immunoblot of TP expression in PBMCs. PBMCs were sorted according to their expression of CD3 (T lymphocytes), CD20 (B lymphocytes) and CD14 (monocytes). Representative of three independent experiments. Total protein amount corresponds to 15µg. (B) Immunoblot of TP expression in CD14^-^ PBMC, CD14^+^ PBMC and monocyte derived macrophages. Protein amount corresponds to 20 000 cells.

**Supplemental Figure 5.**
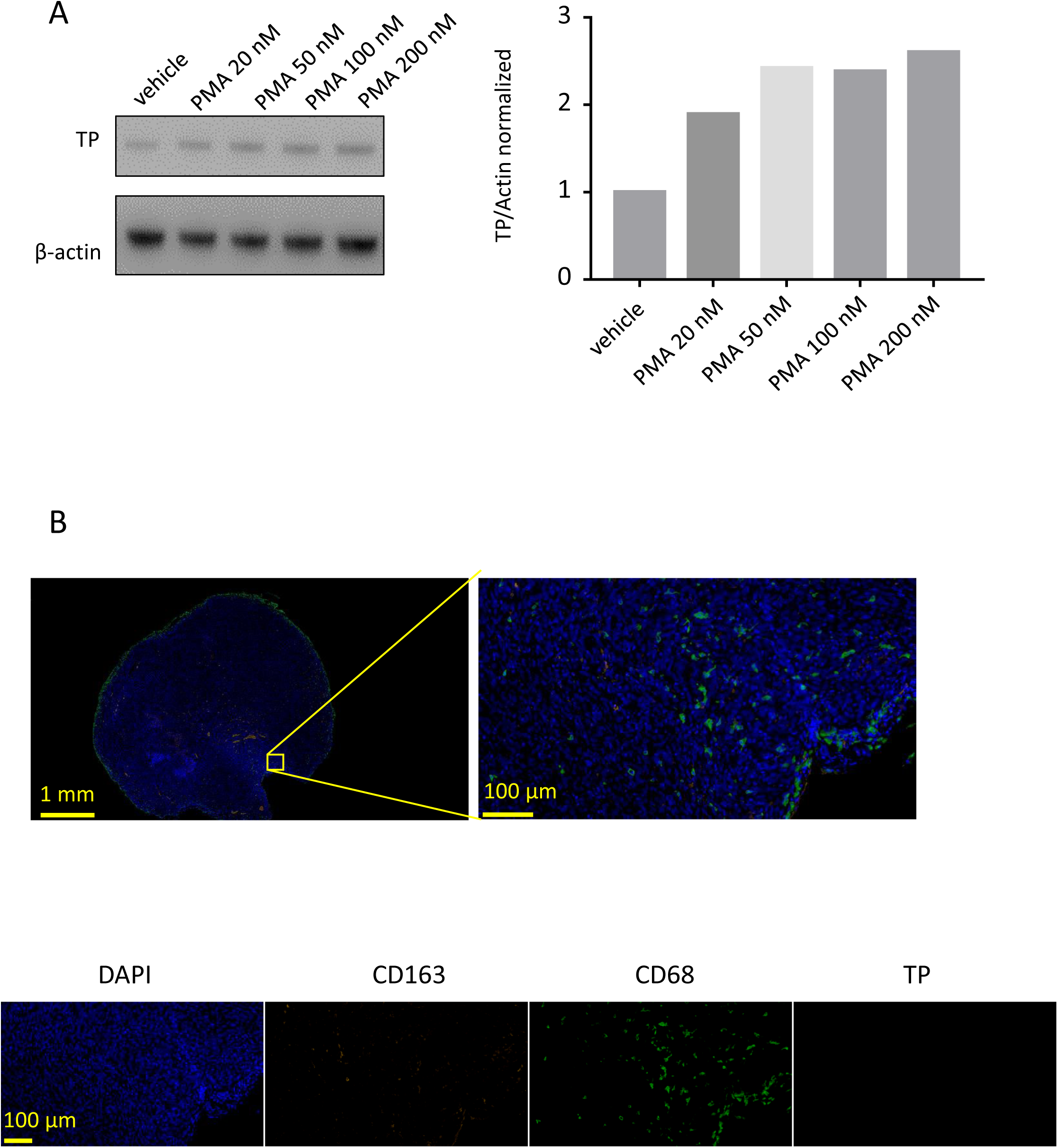
(A) Immunoblot analysis of TP expression in THP1 and THP1 differentiated macrophages with increasing dose of PMA. (B) Immunofluorescence analysis of TP, CD68, CD163 expression in tumor tissue in mice without human macrophage implantation. Nuclei were stained with DAPI.

**Table S1:** Clinical information.

| Patients | Histologic diagnosis | Surgery | TNM | Molecular characterization |
| --- | --- | --- | --- | --- |
| Adult Patient 1 | Adenocarcinoma | Right colectomy (coelioscopic) | pT4N1R0 | p-MMR |
| Adult Patient 2 | Adenocarcinoma poorly differentiated | Right colectomy (coelioscopic) | pT3N1R0 | MSI (loss MSH6) |
| Adult Patient 3 | Adenocarcinoma poorly differentiated low grade | Sigmoidectomy | pT3N2bR0 | NA |
| Adult Patient 4 | Adenocarcinoma well differentiated on tubulovillous adenoma | Left Colectomy | pT3pN1R0 | pMMR |
| Adult Patient 5 | Adenocarcinoma (lieberkühnien) | Left Colectomy | pT3N0 | MSS |
| Adult Patient 6 | Adenocarcinoma (lieberkühnien) low grade | Left Colectomy | pT3N0M0 | MMR |
| Adult Patient 7 | Adenocarcinoma low grade | Right colectomy (coelioscopic) | pT4N1b | MMR |
| Adult Patient 8 | Adenocarcinoma (lieberkühnien) low grade | Right colectomy (coelioscopic) | pT3N0R0 | NA |

